# Nitric oxide inhibits platelet adhesion to platelet-microparticles through reducing integrin α_IIb_β_3_ activation

**DOI:** 10.64898/2026.07.29.741401

**Authors:** Deanna Howley, James Salt, Sam Hall, Matthew S Hindle, Jim Boyne, Wayne Roberts

## Abstract

Increased platelet microparticle (PMP) levels in individuals with risk factors for cardiovascular disease correlate with clinical outcomes in these patient groups. PMPs promote thrombosis through enhancing platelet aggregation and binding to the sub-endothelial matrix following vascular injury. Thus, PMPs behave as soluble ligands and adhesive substrates for platelets, and may drive cardiovascular disease progression. Nitric oxide (NO) is released continually from the endothelium as a potent regulator of platelet activation that is crucial to the balance between haemostasis and thrombosis. However, it is unknown if NO regulates PMP-induced platelet activation.

In this study we isolated platelets and PMPs from whole blood and measured their interactions in adhesion assays and by flow cytometry. Platelet activation was analysed by ELISA for ADP and thromboxane-B2; both secondary platelet agonists released by activated platelets which enhance thrombosis. The affinity upregulation of the principal platelet integrin receptor responsible for platelet aggregation, integrin α_IIb_β_3_, was measured using the antibody PAC-1. Our data show that PMP induced platelet adhesion was associated with, and partially dependent upon, platelet ADP release, TxA2 production and α_IIb_β_3_ upregulation. Crucially, NO dose-dependently reduced these events through cGMP dependent signalling. This is the first report that NO signalling can regulate PMP induced platelet activation and may open an avenue of exploration for clinically targeting PMP driven cardiovascular disease processes.

## Introduction

Platelet activation at sites of atherosclerotic plaque rupture is the leading cause of acute coronary syndromes(1), resulting in over 150,000 hospitalisations and a significant burden of mortality within the UK each year. The advancement of atherothrombosis by platelets occurs in part through their propensity to release platelet microparticles (PMPs). PMPs are small membrane bound vesicles, 0.1-1μm in diameter, shed from activated or apoptotic platelets that account for 70-90% of all circulating microparticles(2). PMPs play a significant role in vascular injury and inflammation through altering endothelial cell cytokine release, enhancing monocyte recruitment to inflamed vessels and by regulating angiogenesis (1, 3, 4). PMPs levels are increased by the presence of cardiovascular risk factors including obesity (5), type 2 diabetes mellitus (6) and progression of atherosclerosis (7), and elevated circulating PMPs are associated an increased clinical risk of cardiovascular disease (8).

PMPs additionally promote thrombosis through enhancing platelet aggregation (9), increasing platelet binding to the endothelium and to the sub-endothelial matrix (10, 11), and by providing a pro-coagulant surface for fibrin formation (12). PMPs are additionally internalised by endothelial cells, inducing a pro-adhesive phenotype through promoting the translocation of von-Willebrand factor (vWF) from Weibel-Pallade bodies to the membrane surface (13). Crucially, PMPs accelerate platelet and fibrin deposition to atherosclerotic arterial walls, enhancing atherothrombosis (14). This places PMPs in the unique position of acting as both an adhesive substrate for platelet deposition and as a circulating direct platelet activator(15).

Nitric oxide (NO), released continually from endothelial cells in response to pulsatile blood flow, maintains a non-thrombogenic barrier within the vasculature. NO is a potent regulator of platelets; inhibiting platelet adhesion, (16–19) aggregation and shape change (20). Once released from the endothelium, NO rapidly diffuses across platelet membranes and binds to soluble guanylyl cyclase (sGC), leading to production of cyclic guanosine monophosphate (cGMP) and subsequent activation of protein kinase G (PKG) (21). PKG phosphorylates a range of intracellular proteins within platelets, limiting activation (22). NO additionally regulates platelet function through cGMP independent S-nitrosylation events (23). Crucially however, it is unknown if PMP induced platelet activation is regulated by NO. Given the importance of platelet activation by PMPs in atherosclerosis (2) and the critical cardio-protective role of NO (24), we investigated the ability of NO to dampen PMP mediated platelet adhesion and activation.

Our data show for the first time that platelet activation by, and adhesion to, PMPs is dependent on the platelet released soluble agonists ADP and thromboxane A2 (TxA2) and the primary platelet integrin α_IIb_β_3_. Furthermore, we demonstrate that NO-cGMP signalling regulates platelet activation by PMPs through reducing α_IIb_β_3_ activation driven by TxA2 and ADP release.

## Materials and methods

### Platelet isolation

Human blood was taken by venepuncture into acid citrate dextrose vacutainers (BD, 366645) by consent from healthy adult donors. All human work was approved by Leeds Beckett University Local Research Ethics Co-ordinator (125755). Donor recruitment started in 01/03/2024, and ended on 19/02/2026. Platelet-rich plasma (PRP) was obtained by centrifugation of whole blood at 200 × *g* at 20 °C, 20 min. Platelets were isolated from the PRP by centrifugation at 800 × *g* at 20 °C, 12 min in the presence of prostaglandin E_1_ (PGE_1_; 50 ng mL^−1^). For some experiments whole blood was incubated with DiOC6 (1µM) followed by platelet isolation. Platelets were resuspended in Modified Tyrodes buffer (150 mmol L^−1^ NaCl, 5 mmol L^−1^ HEPES, 0.55 mmol L^−1^ NaH_2_PO_4_, 7 mmol L^−1^ NaHCO_3_, 2.7 mmol L^−1^ KCl, 0.5 mmol L^−1^ MgCl_2_, 2 mmol L-1 Calcium, 5.6 mmol L^−1^ glucose). For DiO6 stained microparticles, PRP was incubated with 10 μmol DiO6 dye for 15 minutes before the isolation of platelets.

### PMP isolation

PMPs were isolated as we previously reported(25). Briefly, platelet suspensions at 1 × 10^9/mL platelets were activated with thrombin (0.1 U/ml) for 60 minutes at 37 °C. Platelet activation was stopped by EDTA (20mM) and PGE1 (50 ng/mL), and platelets were pelleted by centrifugation (1600 x g; 12 minutes, 3200 x g;15 minutes). PMPs were isolated by centrifugation of platelet free supernatant at 20,000 x g; 90 minutes at 18 °C, then resuspended in Modified Tyrode’s buffer. PMPs were analysed by flow cytometry to confirm expression of a known platelet marker (CD41a) and phosphatidylserine. Flow cytometric analysis was conducted using a BD Accuri C6 Plus Flow cytometer, using a 488 nm laser. To confirm the size of the PMP population (stated in the literature as between 100 – 1000 nm (2)), Nanoparticle Tracking Analysis (NTA) was used (NS300, Malvern Panalytical). The measurements were processed by NTA software, with the size distribution profile data from 3 x 90 second video’s per sample being shown over-plotted.

### Platelet adhesion assay

Ninety-six-well microtiter plates were coated with 100μL of PMPs in suspension (100μg mL−1), for 24 h at 4°C, and blocked at room temperature using 5% BSA diluted in PBS. DioC6 labelled platelets were incubated with indicated inhibitors. Subsequently, platelets (1×10^8^ platelets mL−1) were added to each well and adhered at 37°C for the indicated time. After removal of non-adherent platelets by two PBS washes, adherent platelets were analysed by reading fluorescent signal at excitation-emission 484/501 nm.

### Platelet-PMP binding

For platelet-PMP binding experiments, 100µg/mL DiOC6 stained PMPs were incubated with PerCP-conjugated anti-CD41a stained platelets (3 × 10^7^ cells/mL), in the presence or absence of inhibitors, for 30 minutes. Samples were immediately fixed with 1% paraformaldehyde (PFA) following incubation. Fixed samples were analysed immediately following fixation using flow cytometry (BD Accuri™ C6 Plus). For each sample, 10,000 single platelet events were collected, and results presented as the mean fluorescent intensity of DiOC6 as an approximation of the quantity of PMPs bound to platelets.

### Measurement of ADP and TxA_2_ levels

Platelets (2 × 10^8^ per mL) were pre-incubated with various inhibitors and GSNO before being allowed to adhere to PMPs (100 μg ml) for 0-60 min. TxB_2_ or ADP levels were assayed with an enzyme immunoassay system from Abcam (ab133022, ab83359).

### PAC-1 binding assay

Platelets (1 × 10^8^ mL^−1^) incubated with the appropriate inhibitors were allowed to adhere to six-well microtiter plates coated with PMPs (100 μg mL) for 60 min at 37 °C. Non-adherent platelets were removed by washing with PBS. PAC-1 binding was analysed as we previously reported (19). Data are shown as absorbance values at 405nm.

#### Data sharing

Data is available from the authors on request

### Statistical analysis

Results are expressed as mean ± SEM and were analysed using the Student’s *t*-test or Anova with tukeys post-hoc analysis as stated. All assays were carried out in a minimum of triplicate and with independent platelet donors. The results were considered significant when *P* values were < 0.05.

## Results

### PMPs support platelet adhesion

Initially we confirmed that our PMP isolation produced microparticles of the expected phenotype before examining the ability of PMPs to both bind to platelets in solution and to support platelet adhesion when immobilised. NTA and flow cytometric analysis demonstrated PMPs were 100-1000nm in size (Fig 1A & B), with 93% expressing the phosphatidylserine and the platelet marker CD41A (Fig 1C & D). Furthermore, washed platelets bound to the PMPs whether the microparticles were in suspension (Fig 2B) or immobilised (Fig 2A) as an adhesive ligand. Robust binding occurring at the earliest time point tested (15 minutes) (Fig 2) and was maintained for the longest time point tested (60min).

**Fig 1.**
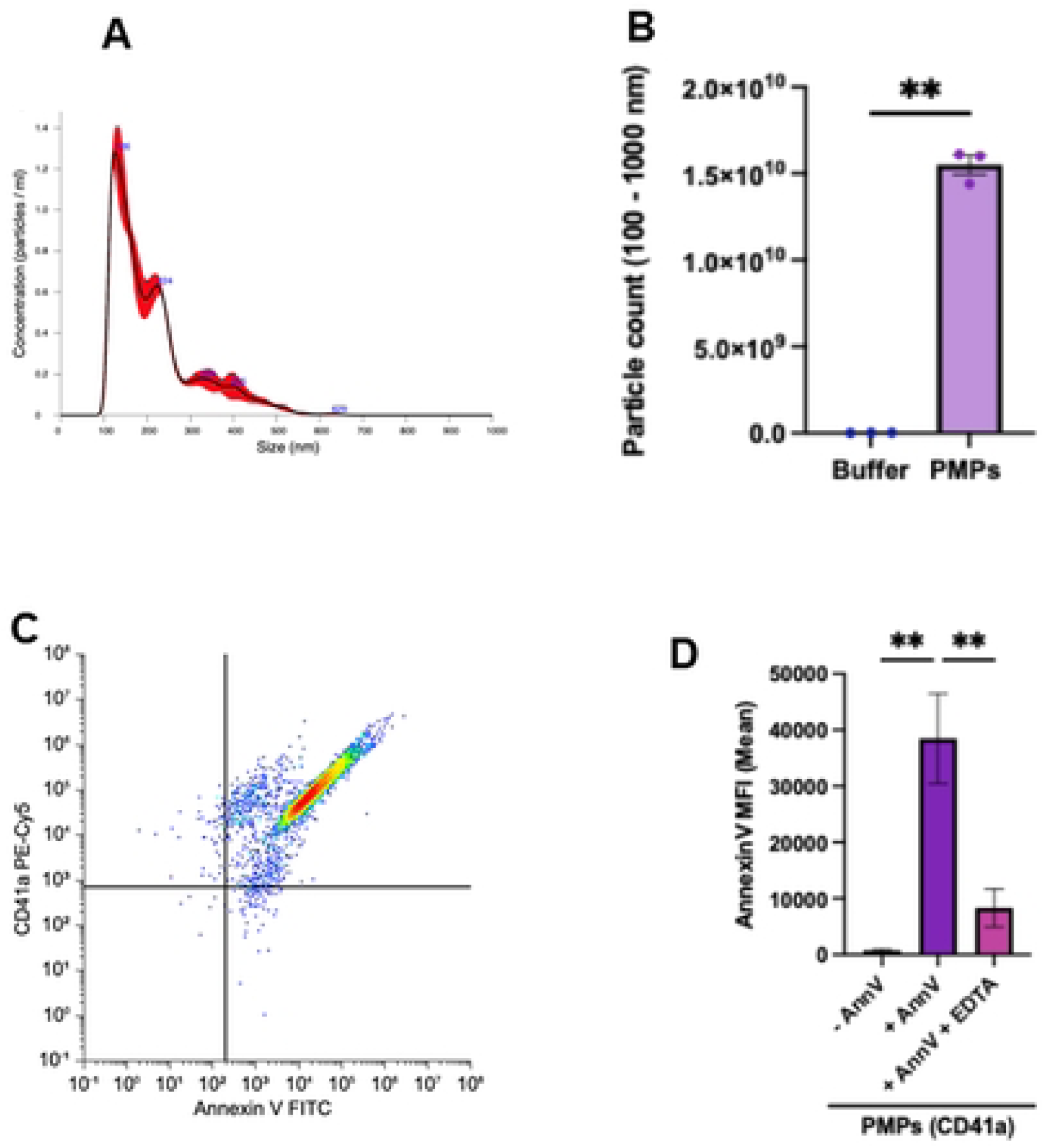
Characterisation of PMPs. Platelets (1×10^9^/mL) were stimulated with thrombin 1U/ml for 60 min at 37°C and PMPs isolated by ultracentrifugation. (A) Representative NTA trace (B) analysis showing average PMP count and size. (C) representative trace and (D) mean values, showing Annexin V (FITC) and CD41a (Pe Cy5) staining. Data are the mean ± SEM of three independent experiments.

**Fig 2.**
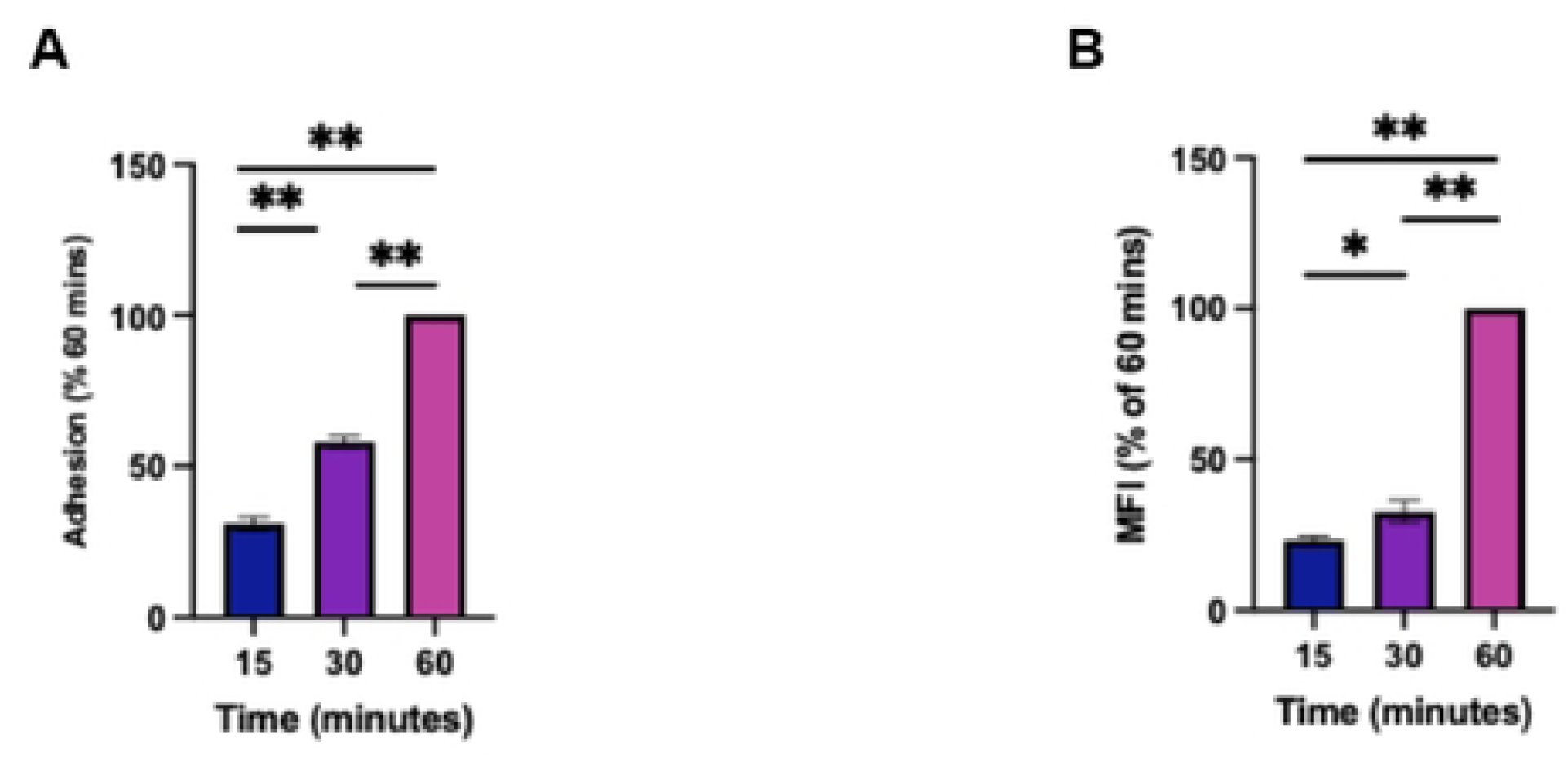
Platelets bind to PMPs in a time dependent manner. (A) Platelets (1×10^8^/mL) were allowed to adhere to 96-well plates coated with PMPS (100/μg mL) for 15-60 min at 37°C. Data are shown as percent adhesion compared with platelets adherent at 60mins. (B) 100µg/mL DiOC6 stained PMPs were incubated with PerCP-conjugated anti-CD41a stained platelets (3 × 10^7^cells/mL), for 15-60 minutes. Samples were fixed with 1% paraformaldehyde (PFA) following incubation. Fixed samples were analysed by flow cytometry (BD Accuri™ C6 Plus). Results are presented as the % mean fluorescent intensity compared to 60 min binding. Values are the mean ± SEM of three independent experiments. Data was analysed by one-way ANOVA and Tukeys *P<0.05, **P<0.001

### Platelet binding and adhesion to PMPs requires the released soluble agonist ADP and TxA2 in addition to the integrin αIIbβ3

Having confirmed that PMPs bind to platelets and induce platelet adhesion, suggesting PMPs promoted platelet activation, we explored underlying mechanisms. Integrin α_IIb_β_3_ is the most highly expressed receptor on platelets with a copy number of 50,000-100,000 (26). Integrin α_IIb_β_3_ is initially present on platelets in a closed confirmation and needs inside out signalling to expose the ligand binding domain. The signalling driving integrin activation and subsequent adhesion to it primary ligand fibrinogen is strongly reliant on the platelet released soluble agonists ADP and TxA2 (27). We therefore examined if PMPs induced α_IIb_β_3_ activation on platelets and if they caused platelet release of ADP and TxA2. Using PAC-1, an antibody specific for activated α_IIb_β_3_, we show sustained activation of α_IIb_β_3_ on platelets attached to PMPs (Fig 3C). Following a similar time course, platelets adherent to PMPs released both ADP and TxA2 (measured as the stable metabolite TxB2). Both agonists were detectable as early as 15 minute after adhesion, with maximum levels being achieved at 60mins (Fig 3), although peak TxB2 production was less than previously observed with collagen stimulation (28), suggesting PMPs alone are a moderate agonist. We next examined if these events are required for platelet adhesion to PMPs. Treating platelets with the α_IIb_β_3_ inhibitory peptide RGDS significantly reduced both platelet-PMP binding in suspension and PMP induced adhesion by 16±4% and 33±9% respectively (P<0.05) (Fig 4A & B). Inhibiting platelet derived ADP and TxA2 with apyrase and indomethacin, similarly reduced platelet adhesion and binding in suspension (Fig 4A & B). These data show that activated integrin α_IIb_β_3_ and platelet derived ADP and TxA2 and required for maximum binding and adhesion to PMPs. Importantly the addition of RGDS to apyrase and indomethacin treated platelets had no further additive inhibitory effect (Fig 4). These data suggest that ADP, TxA2 and α_IIb_β_3_ enhance binding through a similar co-dependent mechanism. We therefore assessed if integrin α_IIb_β_3_ activation on platelets adherent to PMPs was dependent on released ADP and TxA2. Apyrase and indomethacin treatment of platelets prior to incubation with PMPs significantly reduced the active form of the integrin (P<0.05). Together these data suggest platelet attachment to PMPs induces the release of ADP and TxA2 from platelets, which together help upregulate the affinity of integrin α_IIb_β_3_, enhancing firm adhesion.

**Fig 3.**
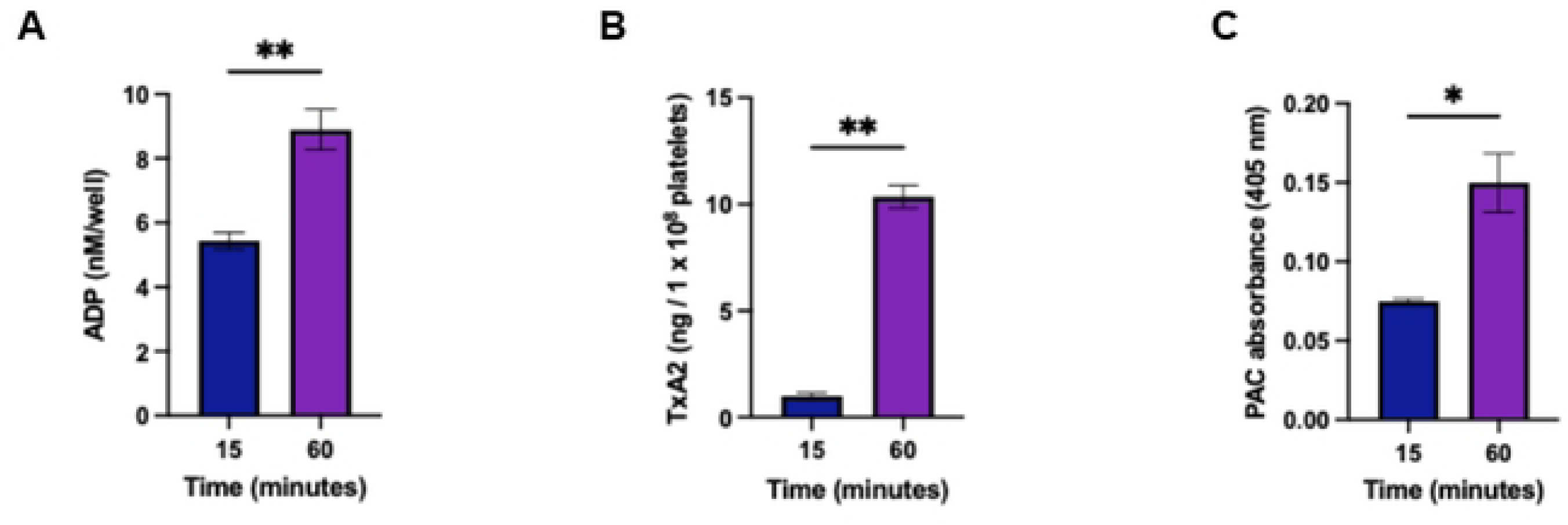
Platelet adhesion to PMPs induces ADP release, TxA2 secretion and integrin α_IIb_β_3_. Platelets 2×10^8^ ml were allowed to adhere to immobilised PMPs (100µg/ml) for 15-60min. ADP release (A), TxB2 release (B) and C) PAC-1 binding (absorbance 405 nm) were measured a15-60min. Data shows is mean of three repeats ±SEM, data was analysed by students t test., (P<0.05, *), (P<0.001,**).

**Fig 4.**
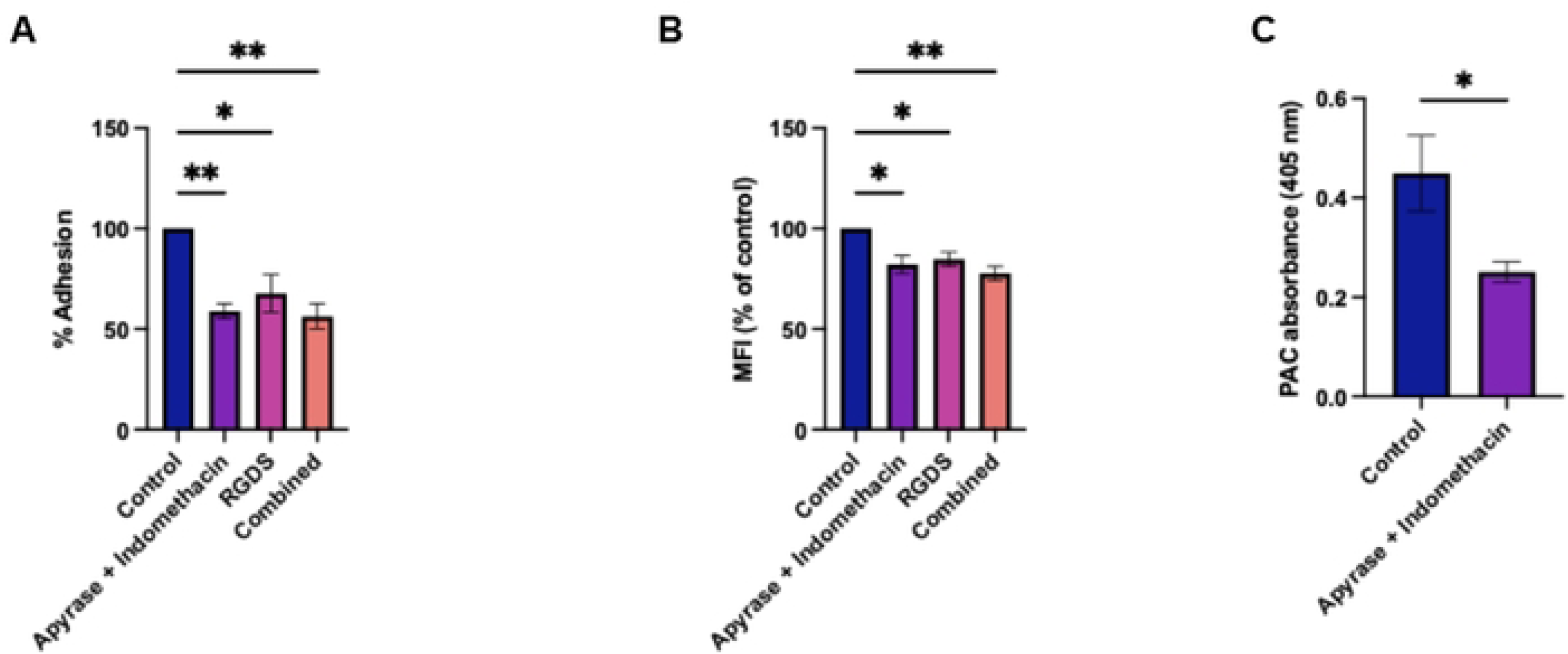
Platelets binding to platelet microparticles is reliant on secondary mediators and the integrin α_IIb_β_3_. (A) Platelets (1×10^8^/mL) were allowed to adhere to 96-well plates coated with PMPS (100/μg mL) for 60 min at 37°C in the presence of combinations of apyrase, indomethacin and RGDS. Data are shown as percent adhesion compared with platelets adherent for 60mins in the absence of inhibitors. (B) 100µg/mL DiOC6 stained PMPs were incubated with PerCP-conjugated anti-CD41a stained platelets (3 × 10^7^ cells/mL), for 60 minutes in the presence of indicated inhibitors. Fixed samples were analysed by flow cytometry (BD Accuri™ C6 Plus). Results are presented as the % mean fluorescent intensity compared to 60 min binding in the absence of inhibitors. (C) Platelets (2×10^8^/mL) were allowed to adhere PMPS (100/μg mL) for 60 min at 37°C in the presence of indicated inhibitors. PAC-1 binding is show as absorbance at 405nm. Values are the mean ± SEM of three independent experiments. Data was analysed by one-way ANOVA and Tukeys or students t test *P<0.05, **P<0.001

### Nitric oxide reduces platelet adhesion to PMPs

NO is a potent platelet inhibitor that we have previously shown reduces platelet adhesion to the extracellular matrix proteins collagen (18), fibrinogen (16) and von-Willebrand factor (vWF) (19). Given data that PMPs support platelet adhesion and activation (9, 11, 14, 29, 30) and the suggested role PMP induced platelet activation plays in vascular disease (1), we sought to determine if NO could reduce platelet binding and adhesion to PMPs. Treating platelets with the NO donor GSNO (0-100μM) reduced platelet adhesion to PMPs in a dose dependent manner. The lowest dose of GSNO tested (1μM) had no significant effect on adhesion or PMP binding, by 100µM GSNO inhibition of adhesion reached 31±9% (Fig 5A) (P<0.05). GSNO was equally effective at reducing PMP-platelet binding under stirring conditions, mimicking interactions that happen in flowing blood (Fig 5B). Having confirmed the inhibitory effect of GSNO on PMP-platelet interactions, we examined the underpinning signalling. NO regulates platelet function through cGMP dependent and independent mechanisms (21). To differentiate between the pathways we used ODQ which is a potent and selective inhibitor of the cGMP producing enzyme sGC, as previously reported (18). GSNO induced inhibition of platelet adhesion was reduced from 20±8% to 0±4% by ODQ (p<0.05) and in suspension from 30±6% to 10±4%. These data suggest that maximum inhibition of platelet binding to PMPs by NO requires cGMP dependent signalling pathways. To confirm these findings, we used the cGMP mimetic 8-Br-cGMP. Direct activation of the cGMP-PKG pathway with 8-Br-cGMP reduced platelet adhesion to PMPs by 19±5% and binding in suspension by 25±8%, not significantly different that induced by GSNO, confirming the major role played by cGMP dependent signalling (Figure 5C & D).

**Fig 5.**
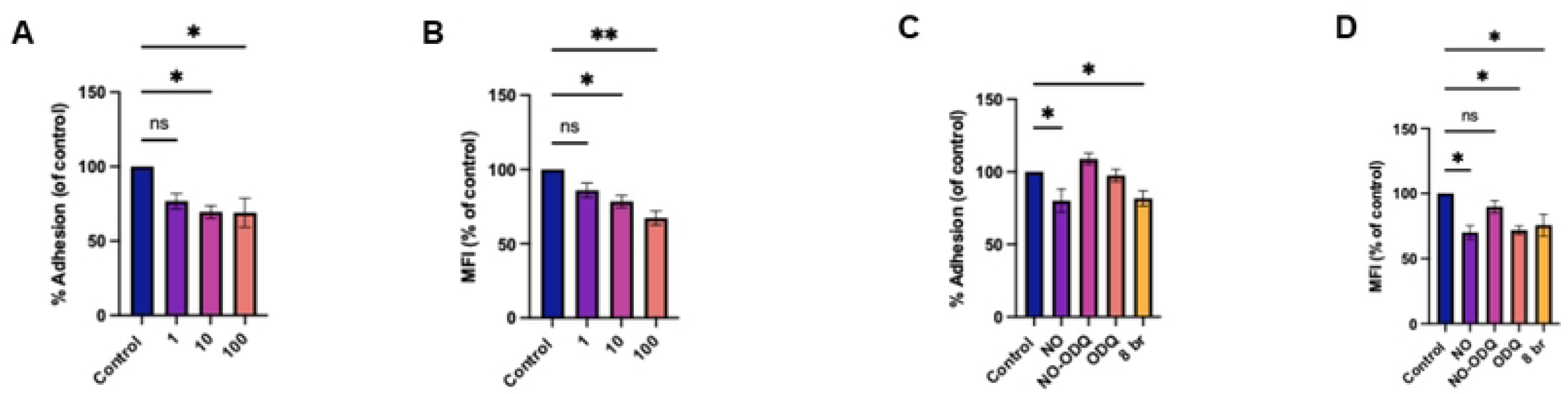
Platelets binding to platelet microparticles is inhibited by NO. (A & C) Platelets (1×10^8^/mL) were allowed to adhere to 96-well plates coated with PMPS (100/μg mL) for 60 min at 37°C in the presence of GSNO (0-100µM) and indicated inhibitors. Data are shown as percent adhesion compared with platelets adherent for 60mins in the absence of inhibitors. (B & D) 100µg/mL DiOC6 stained PMPs were incubated with PerCP-conjugated anti-CD41a stained platelets (3 × 10^7^ cells/mL), for 60 minutes in the presence of GSNO and indicated inhibitors. Fixed samples were analysed by flow cytometry 60 minutes C6 Plus). Results are presented as the % mean fluorescent intensity compared to 60 min binding in the absence of inhibitors. Values are the mean ± SEM of three independent experiments. Data was analysed by one-way ANOVA and Tukeys *P<0.05, **P<0.001

### NO regulates platelet adhesion to PMPs through targeting integrin α_IIb_β_3_ and secondary mediator driven adhesion

Having confirmed the inhibitory effect of NO on platelet adhesion and binding to PMPs we examined potential targets. Our data thus far show that platelet interaction with PMPs has a component reliant on secreted agonist induced activation of integrin α_IIb_β_3_, as well as component not reliant upon these mechanisms. To examine which of these was targeted by NO we measured adhesion and binding in the presence of RGDS, apyrase and indomethacin plus minus GSNO. If NO reduced binding when the integrin and secondary mediators were pharmacologically inhibited it would suggest a target distinct from these pathways. The data show combined inhibition of integrin α_IIb_β_3_, ADP and TxA2 or treatment with GSNO all reduced adhesion to PMPs by the same extent. Importantly, combined treatment of RGDS, apyrase and indomethacin with GSNO had no further additive effect on inhibition of adhesion compared to the inhibitors without GSNO (Fig 6A). Blocking the integrin and secondary mediators reduced adhesion by 49±6% which was 47±6% when GSNO was used in addition. These data suggest NO targets the secondary mediator driven α_IIb_β_3_ component of platelet adhesion to PMPs. Whilst the pattern was similar in suspended platelets, there was also a degree of additive inhibition. These data suggest NO may have additional distinct targets when regulating platelet-PMP binding in suspension, but not when PMPs were acting as an immobilised adherent surface. We therefore investigated if NO could directly reduce ADP and TxA2 release and activation of integrin α_IIb_β_3_.

**Fig 6.**
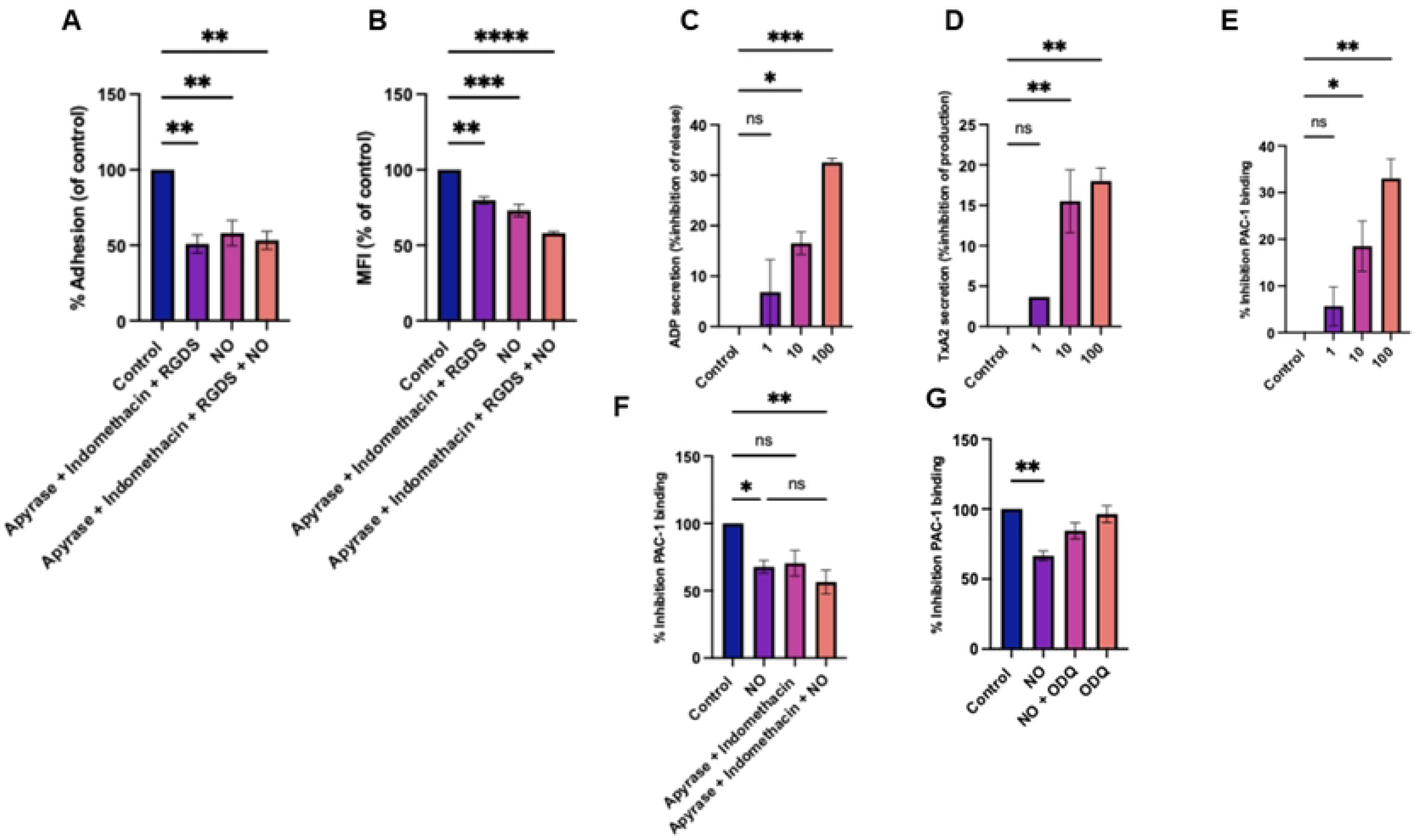
NO regulation of platelet binding to platelet microparticles occurs through targeting α_IIb_β_3_. (A) Platelets (1×10^8^ /mL) were allowed to adhere to 96-well plates coated with PMPS (100/μg mL) for 60 min at 37°C in the presence of GSNO and indicated inhibitors. Data are shown as percent adhesion compared with platelets adherent for 60mins in the absence of inhibitors. (B) 100µg/mL DiOC6 stained PMPs were incubated with PerCP-conjugated anti-CD41a stained platelets (3 × 10^7^ cells/mL), for 60 minutes in the presence of GSNO and indicated inhibitors. Fixed samples were analysed by flow cytometry (BD Accuri™ C6 Plus). Results are presented as the % mean fluorescent intensity compared to 60 min binding in the absence of inhibitors. Platelets 2×10^8^ ml were allowed to adhere to immobilized 100µg/ml PMPs for 60min in the presence of GSNO (0-100µM). ADP release (C), TxB2 release (D) and (E-G) PAC-1 binding (absorbance 405 nm) were analysed. Values are the mean ± SEM of three independent experiments. Data was analysed by one-way ANOVA and tukeys *P<0.05, **P<0.001

### NO inhibits PMPs induced αIIbβ3 activation through targeting secondary mediator release

Treating platelets adherent to PMPs with GSNO had a small but significant inhibitory effect on TxA2 production and a more pronounced impact on ADP release. GSNO (1µM) reduced TxA2 release by 3.6±0.5% which increased to 17.7±2% in the presence of GSNO (100µM, Fig 6D). Inhibition of ADP release ranged from 7±2% to 33±0.5 as the concentration of GSNO was increased. Importantly GSNO also dose dependently reduced integrin α_IIb_β_3_ activation as assessed by PAC-1 binding (Fig 6E). 1µM GSNO reducing PAC-1 binding by 6±3% whilst 100µM reducing binding by 33.5±4 % (P<0.05). These data show that whilst NO reduces PMP induced platelet secondary mediator release and integrin α_IIb_β_3_ activation, a proportion of these pathways are refractory to the effects of NO, in line with our data that NO cannot fully abolish platelet adhesion to PMPs. To investigate if NO was targeting secondary mediator dependent or independent integrin activation we measured PAC-1 binding, as a marker of integrin α_IIb_β_3_ activation, on platelets adherent to PMPs treated with apyrase and indomethacin in the presence and absence of GSNO. Blocking the effects of ADP and TxA2 reduced α_IIb_β_3_activation by 29.6±7% (p<0.05), incubation with GSNO reduced activation by 32±4% (p<0.05) and combined treatment of apyrase, indomethacin and GSNO reduced activation by 43±7% (p<0.05) which was not significantly different than treatment with GSNO alone (Fig 6F). These data suggest NO principally regulates integrin α_IIb_β_3_ activation and subsequent adhesion through moderating the release of TxA2 and ADP. Finally, to determine how NO targeted integrin α_IIb_β_3_ activation, we used ODQ to inhibit the cGMP pathway (Fig 6). Under these conditions NO mediated inhibition of PAC-1 binding (36±4% p<0.05) was reduced to 15.5±7% (NS vs control), suggesting that NO mediated inhibition of integrin activation occurs through a cGMP dependent pathway.

## Discussion

A wealth of data show that PMPs strongly promote cardiovascular disease progression (1, 31). In part this occurs from the procoagulant nature of PMPs which support the generation of thrombin, driving clot formation. However there is a growing body of evidence that PMPs additionally support direct platelet activation; crucially platelet driven thrombosis is the underlying cause of most acute coronary syndromes (14). A greater understanding of platelet-PMP cross talk, and its regulation by nitric oxide (NO); the chief *in vivo* platelet inhibitor, is therefore needed to further our understanding of thrombosis. In this study we report for the first time that NO significantly reduced both platelet binding to PMPs in suspension and to immobilised PMPs. Platelet-PMP interactions in suspension are a proxy model for how these interactions might occur in flowing blood; a scenario important in CVD patients where platelets are presented partially activated and hence primed to bind to a range of targets, resulting in enhanced activation. Platelet adhesion meanwhile is the critical first step in thrombosis.

PMPs comprise 60-90% of all circulating microparticles (32) and their levels increase in unstable angina (33), acute myocardial infarction (34) and in atherosclerosis (2, 35). Increased PMP levels correlate with a number of parameters including carotid artery intimal thickness and plaque burden in these conditions (2). The mechanisms linking these observations are myriad, but include enhancing thrombosis, vascular dysfunction and inflammation (15). Conversely, PMP deficiency is associated with a prolonged bleeding time, as seen in patients with Castaman’s defect and Scotts syndrome (2, 36), who also have altered platelet aggregation under arteriolar flow (37). In addition to their procoagulant nature, PMPs interact with components of vessel walls such as collagen, fibrinogen and fibronectin, providing an enhanced substrate for platelet adhesion (11). We confirm these findings, reporting here that PMPs support platelet adhesion in time dependent manner. PMP induced platelet adhesion is likely to be clinically relevant given that PMPs significantly enhance agonist induced platelet aggregation (9), and under flow conditions platelet deposition to damaged porcine arteries and human atherosclerotic plaques is significantly increased in PMP enriched blood (14). Furthermore, using laser induced vascular endothelial injury in mice Falati et al demonstrated that microparticles bind to activated platelets, contributing to thrombus formation (38), and platelet have recently been demonstrated to be able to internalise PMPs, enhancing aggregation and clot formation (30).

Elucidating the molecular mechanisms by which PMPs might evoke thrombus formation is crucial for improving our understanding of their role in atherosclerotic heart disease. Initial platelet adhesion is largely driven by integrin receptors, with released secondary agonists such as ADP and TxA2 then potentiating this response through upregulation of integrin affinity (39). Previous studies have suggested that integrin α_IIb_β_3_ is responsible for mediating PMP-platelet binding (11) although have not directly measured this, or investigated how PMPs activate the integrin. We show that PMPs induce ADP release and TxA2 production from adherent platelets in addition to upregulating the affinity of integrin α_IIb_β_3_ as shown by PAC-1 binding. These responses were needed for maximal platelet adhesion to PMPs, as demonstrated by their respective inhibitors reducing platelet binding. It is likely that released ADP and TxA2 potentiate adhesion to PMPs through indirect activation of integrin α_IIb_β_3_, as incubating platelets with the peptide RGDS to abrogate integrin binding reduced adhesion to the same extent as treatment with apyrase and indomethacin to block ADP and TxA2 respectively, and co-incubation with these compounds had no additive effect. ADP and TxA2 have previously been shown to augment integrin α_2_β_1_ adhesion to collagen and integrin α_IIb_β_3_ binding to fibrinogen in a similar fashion (39). Crucially, we show apyrase and indomethacin blunted PAC-1 binding to PMP adherent platelets, confirming the PMP induced platelet release of ADP and TxA2 is needed for integrin α_IIb_β_3_ activation. Simultaneous blockade of integrin α_IIb_β_3_, ADP and TxA2 did not fully reduced platelet adhesion to PMPs however, suggesting other receptor(s) must also play a role in this process. PMPs have been reported to express a range of additional receptors including GPIb, P-selectin and CD36 (40) and CD36 has been demonstrated to be involved in PMP augmented platelet aggregation (29), raising the possibility that CD36 or other platelet receptors may also help mediate platelet-PMP adhesion.

Whilst there is clear evidence that PMPs initiate and augment platelet activation, we present to the best of our knowledge, the first evidence that NO can target and reduce PMP-mediated platelet adhesion and activation. Similar findings have previously been reported when studying the impact of NO on platelet-ECM interactions (18, 19). These data fit with the concept that NO is produced to prevent excessive thrombus formation that could lead to vessel occlusion, rather than fully block platelet activation, which is necessary for haemostasis. Using the sGC antagonist ODQ, under conditions we previously reported to fully block cGMP-PKG signalling (18, 19, 41), significantly reduced the ability of NO to inhibit platelet adhesion to PMPs. These data suggest that NO primarily regulates PMP-platelet binding through cGMP dependent mechanisms. Similarly, using the direct PKG activator 8-Br-cGMP reduced platelet adhesion and binding to PMPs, adding further evidence that cGMP-PKG signalling is able to inhibit platelet activation by PMPs. We cannot however rule out non-cGMP effects, as ODQ did not fully block the effects of NO on platelet binding to PMPs in suspension. Indeed, previous work has shown that NO can cause s-nitrosylation of N-ethylmaleimide-sensitive factor, an ATPase required for degranulation(42) and the integrin α_IIb_β_3_ (43). These mechanisms may explain the residual inhibition of platelet-PMP binding in suspension elicited by NO in the presence of ODQ that we report here.

Together our data show PMP induced platelet adhesion and activation is primarily driven by integrin α_IIb_β_3_. When the integrin was inhibited, the residual adhesion remaining was not reduced by NO treatment, suggesting that NO targets integrin α_IIb_β_3_ activation to reduce PMP-mediated platelet activation. Congruently we show that NO reduced PAC-1 binding in an ODQ sensitive manner, confirming NO-cGMP signalling targets PMP evoked α_IIb_β_3_ activation. The further removal of ADP and TxA2 from this system had no significant additional effect, suggesting NO reduces secondary mediator bioavailability, thereby dampening α_IIb_β_3_ activation and subsequent adhesion. Crucially we demonstrate that NO limits PMP induced platelet release of both ADP and TxA2. Given the well documented role PMP-induced platelet activation plays in cardiovascular disease, further studies are warranted to assess how this can be clinically targeted.

## Acknowledgements

This work was supported by funding from the Centre for Biomedical Science Research, Leeds Beckett University.

## Authorship

Contributions: D.H, J.S and SH performed experiments and analysed data. WR, JRB and MSH. co-designed the study and wrote the manuscript together. All authors have read and agreed to the published version of the manuscript

The authors declare no conflicts of interest.

